# peakScout – a user-friendly and reversible peak-to-gene translator for genomic peak calling results

**DOI:** 10.1101/2025.09.07.671934

**Authors:** Alexander L. Lin, Lana A. Cartailler, Jean-Philippe Cartailler

## Abstract

**Summary:** peakScout is a command line and web-based bioinformatics tool designed to quickly and easily bridge the gap between genomic peak data and gene annotations, enabling researchers to understand the relationship between measurements of regulatory elements and their target genes. At its core, peakScout processes genomic peak files obtained through various means chromatin profiling and maps them to nearby genes using reference genome annotations. The workflow begins with input processing, where peak files are standardized and reference GTF files are decomposed into chromosome-specific feature collections. The core analysis modules then perform bidirectional mapping: peak-to-gene identifies which genes are potentially regulated by specific genomic regions, while gene-to-peak reveals which regulatory elements might influence particular genes of interest. Throughout this process, nearest-feature detection algorithms handle the complex spatial relationships between genomic elements, considering factors like distance constraints and feature overlaps. Finally, the results are formatted into researcher-friendly CSV and Excel outputs, providing a comprehensive view of the genomic landscape that connects regulatory elements to their potential gene targets.

**Availability and implementation:** The web version of peakScout is available at https://vandydata.github.io/peakScout/. The command line version is available at https://github.com/vandydata/peakScout and archived on Zenodo (URL to be provided upon version 1.0 release) under the GNU Affero General Public License v3.0. Installation instructions, example datasets, and detailed usage examples are provided in the GitHub repository README file. peakScout is implemented in Python and is platform independent, but the web version is implemented in Amazon Web Services and thus uses proprietary infrastructure.

## 1 Introduction

High-throughput genomic techniques have revolutionized our understanding of protein-DNA interactions, chromatin modifications, and gene regulation. Methods such as ChIP-seq, CUT&RUN, CUT&TAG, ATAC-seq, DNase-seq, and FAIRE-seq typically generate thousands of genomic regions of interest (peaks) that represent binding sites, accessible chromatin regions, or enriched chromatin marks. However, translating these genomic coordinates into biologically meaningful insights remains challenging, particularly for researchers without extensive bioinformatics expertise.

The critical step of associating genomic peaks with nearby genes is essential for understanding regulatory networks and interpreting experimental results. While several sophisticated tools exist for genomic data analysis, they often require significant computational skills, command-line proficiency, and parameter optimization. BEDTools (Quinlan and Hall, 2010) offers comprehensive functionality for manipulating genomic intervals but demands familiarity with command-line operations and complex parameters. Other tools like HOMER (Heinz *et al*., 2010), GREAT (McLean CY, et al. GREAT improves functional interpretation of cis-regulatory regions - Google Search), and ChIPseeker (Yu *et al*., 2015) provide peak annotation capabilities but either require programming knowledge or offer limited flexibility in defining proximity relationships.

peakScout addresses this gap by providing a straightforward, accessible solution for bidirectional mapping between genomic peaks and genes. Unlike existing tools that may overwhelm users with complex options or require extensive bioinformatics training, peakScout offers an intuitive approach that accepts standard output from popular peak callers like MACS2 (Zhang *et al*., 2008) and SEACR (Meers *et al*., 2019) and produces researcher-friendly results in familiar formats (CSV, Excel). The tool supports both peak-to-gene and gene-to-peak analyses, allowing researchers to either identify potential target genes of regulatory elements or find regulatory elements that might influence specific genes of interest.

By simplifying this critical analytical step, peakScout enables bench scientists and bioinformatics novices to quickly interpret their genomic data without extensive computational training. The tool’s straightforward implementation and accessible output formats facilitate rapid integration with downstream analyses and visualization tools, accelerating the path from raw peak data to biological insights.

## 2 Language and algorithmic approach

peakScout is implemented in Python, leveraging the high-performance Polars data processing library for efficient manipulation of tabular genomic data. This choice of technology enables rapid processing of large genomic datasets while maintaining a user-friendly interface. The core algorithmic approach in peakScout centers around efficient chromosome-specific feature decomposition and nearest-feature detection.

For reference data management, peakScout employs a hierarchical decomposition strategy that organizes genomic features by type (gene, exon, CDS, etc.) and chromosome. This approach, implemented in *decompose_ref*.*py*, significantly reduces the search space when identifying nearest features, as only features on the same chromosome need to be considered. Each feature collection is further organized into separate files sorted by start and end positions, enabling efficient binary search operations when determining proximity relationships.

The nearest-feature detection algorithm employs a bidirectional search strategy that simultaneously evaluates upstream and downstream features while accounting for potential overlaps. This approach is particularly efficient as it leverages the pre-sorted nature of both the decomposed reference data and the input features, allowing for rapid identification of nearest features even for datasets containing thousands of peaks and reference features.

### 2.1 Supported operations

Figure 1 illustrates the general workflow of peakScout. Prior to conducting peak or gene analyses, peakScout implements a decomposition operation that preprocesses reference annotation files – such as GTFs – into a format optimized for downstream analysis. This preprocessing step significantly improves the efficiency of nearest feature identification by partitioning the reference data based on both chromosome and feature type. This decomposition is a one-time operation per reference file, and the resulting data are stored in a user-defined directory for future analyses.

**Figure 1.**
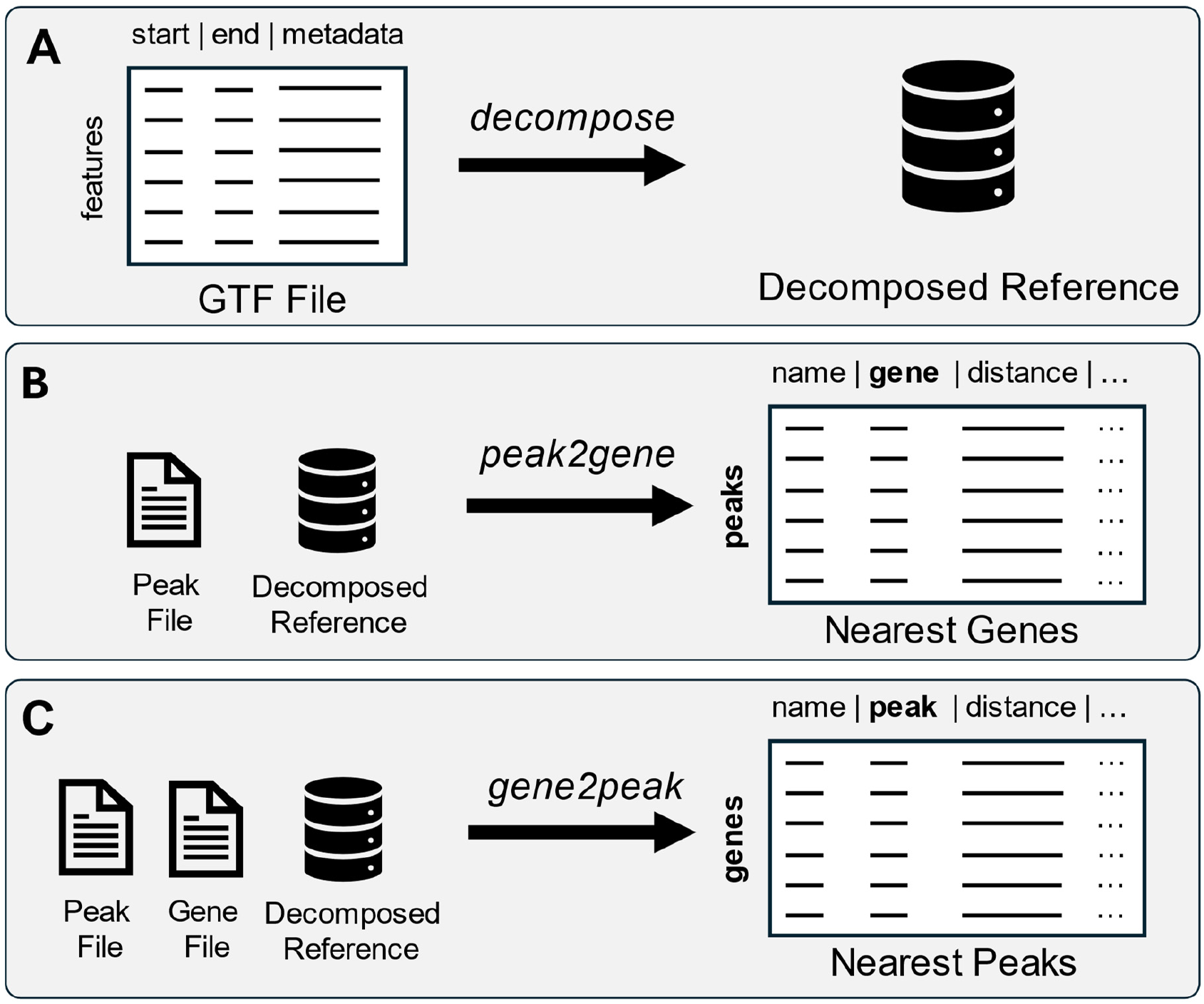
peakScout analysis workflow. The peakScout tool employs a two-step process for bidirectional mapping between genomic peaks and genes. (a) Reference preparation: GTF annotation files containing genomic features (start, end, metadata) are decomposed into chromosome- and feature-specific collections to optimize search efficiency. (b) Peak-to-gene analysis: Input peak files are processed alongside the decomposed reference to identify the k-nearest genes for each peak, generating a comprehensive output table with peak names, associated genes, and distance measurements. (c) Gene-to-peak analysis: A user-provided gene list is analyzed against the peak file and decomposed reference to identify the k-nearest peaks for each gene of interest, producing an output table with gene names, associated peaks, and distance measurements. Both analytical pathways support flexible distance constraints and multiple output formats (CSV, Excel).

Following the decomposition step, users may proceed with one of two analytical functions: *peak2gene* or *gene2peak*. The *peak2gene* function identifies the nearest annotated genes from the reference dataset to a user-defined set of genomic peaks. Conversely, the *gene2peak* function determines the nearest peaks – also provided by the user – relative to a specified list of genes. Both functions provide comprehensive options for defining peak boundaries and proximity constraints. Users can choose between native peak boundaries (as defined by the peak caller), peak summits, or artificial boundaries with user-defined extensions. Additionally, users can specify maximum distance constraints for upstream and downstream features, ensuring that only biologically relevant associations are reported.

For input flexibility, peakScout supports multiple peak file formats through specialized processing functions. Internally defined operations automatically detect and handle MACS2 output (both Excel and BED formats) and SEACR output, standardizing these diverse formats into a consistent internal representation. BED6 is also supported, which provides wide support for most peak-calling outputs. This enables seamless integration with various peak calling workflows without requiring users to perform format conversions. When users want to obtain a list of peaks nearest to their genes of interest, a single-column gene list is all that is required.

To facilitate ease of use, peakScout includes pre-generated reference files for commonly studied organisms: C. elegans (WBcel235), Drosophila (BDGP6.54), zebrafish (GRCz11), S. cerevisiae (R64-1-1), X. tropicalis (v10.1), mouse (mm39 and mm10), human (hg38 and hg19), and pig (Sscrofa11.1). The reference GTF files were downloaded from Ensembl (Dyer *et al*., 2025) and Gencode (Mudge *et al*., 2025), and the resulting decomposed references are available via a public S3 bucket, eliminating the need for users to manually process GTF files for these organisms. Instructions are provided as to how to download and use these pre-generated references.

### 2.4 Feature detection and proximity analysis

The core of peakScout’s analytical capability lies in its feature detection and proximity analysis algorithms. Unlike some tools that simply report the nearest TSS for each peak, peakScout implements a more sophisticated approach that considers the full genomic context.

The *get_nearest_features* function in *process_features*.*py* employs a bidirectional search strategy that simultaneously evaluates upstream and downstream features. This approach allows peakScout to identify the k-nearest features in both directions, providing a more comprehensive view of the genomic neighborhood around each peak. The function also handles feature overlaps, ensuring that genes directly overlapping with peaks are prioritized in the results.

For proximity constraints, peakScout allows users to specify maximum distance thresholds for upstream and downstream features through the *up_bound* and *down_bound* parameters. This functionality enables researchers to focus on biologically relevant associations based on their understanding of regulatory element behavior in their specific experimental context.

The *constrain_features* function further optimizes the search process by filtering reference features based on these distance constraints before performing the detailed proximity analysis. Through binary search operations, it also ensures that, after accounting for potential overlaps, searches for upstream and downstream nearest features begin at the closest possible location. These pre-filtering steps significantly reduce the computational burden for large datasets, enabling rapid analysis even on standard desktop computers.

### 2.5 Output generation and visualization

peakScout provides flexible output options through its *write_output*.*py* module. Results can be exported in both CSV and Excel formats, with the Excel output including additional formatting for improved readability. As shown in Figure 2, the output includes comprehensive information about each peak or gene, including 1) genomic coordinates (chromosome, start, end), 2) feature identifiers (peak name or gene name), 3) k-nearest features (as specified by the user), 4) distance to each nearest feature (with negative values indicating upstream features), and 5) a pre-formed URL to the UCSC genome browser (for supported genomes) that includes a “highlight” of their peak in a rich and interactive genomic context. This rich output format enables researchers to quickly identify potential regulatory relationships and prioritize candidates for further experimental validation.

**Figure 2.**
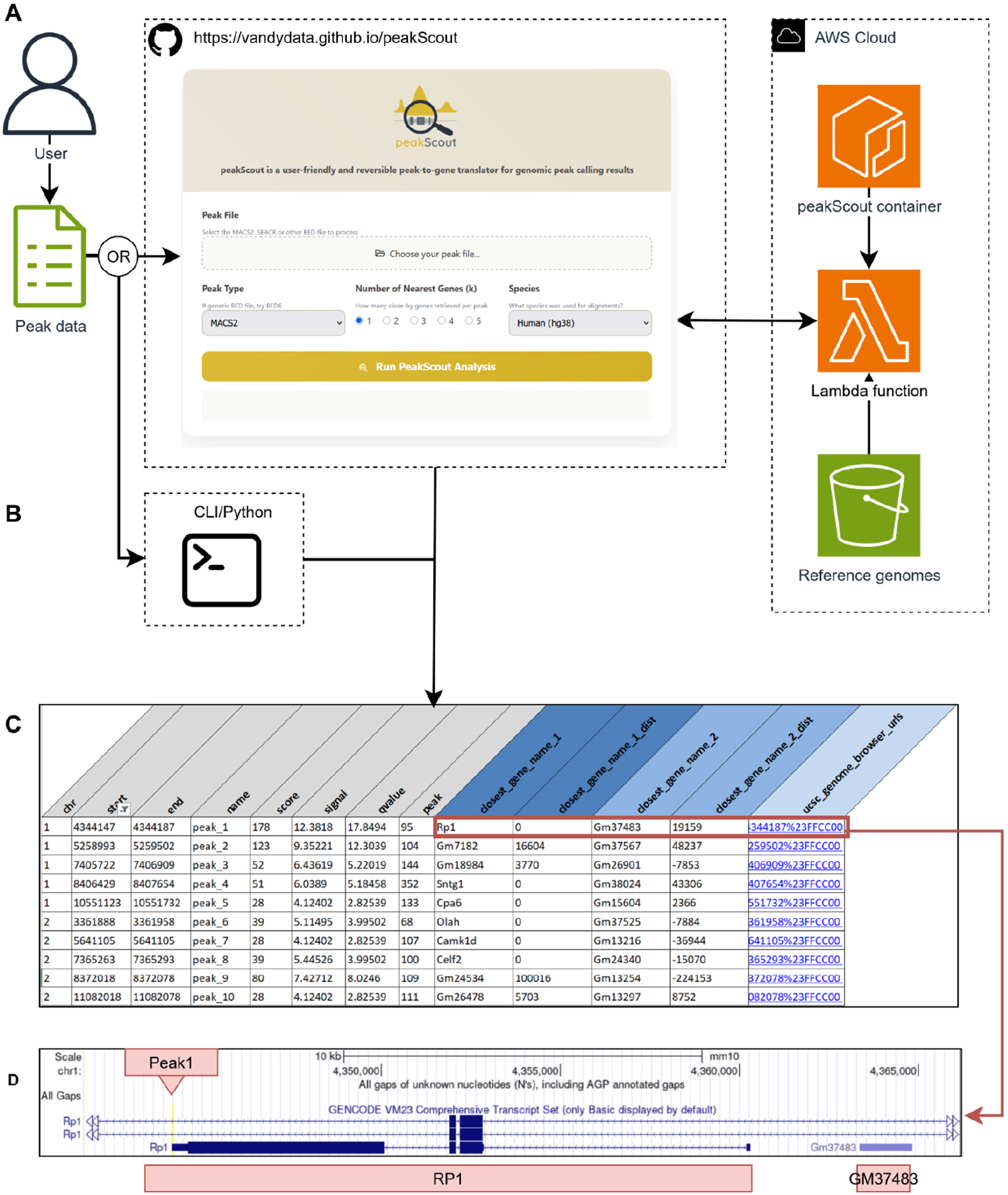
peakScout in practice. A) Users and their peak data can interact with peakScout either via the web version of peakScout at https://vandydata.github.io/peakScout/orB) via the command line. The user will upload a peak file and select basic parameter settings. The result is provided as tabular data, which includes a gene name and genomic distance from the peak, for as many nearby genes selected by the user. For example, in C), the user would have selected 2 nearby genes. For the first peak on chr1:4344147-4344187, the nearest gene is Rp1 (retinitis pigmentosa 1), which is essential of retinal photoreceptor cell function. The second nearest gene is Gm37483. To help with accessing additional gene annotations in this genomic context, a link to the relevant region at the UCSC genome browser is provided, which provides a context specific “highlight” (as per UCSC specifications) of the peak, as shown in D. In this case, the peak is present in the promoter region of Rp1 and most likely plays a role in transcriptional control.

### 2.6 Interfaces

For use as imported functions or at the command line, peakScout can be installed as per the instructions on the GitHub website and dependencies installed using pip or uv, commonly used package managers.

#### 2.6.1. Use as a python library

To use peakScout functions within a typical Python script, one can import the peakScout functions (*peak2gene, gene2peak*, or *decompose_ref*) and use as desired in their program or workflow.

#### 2.6.2. Use via a website and serverless cloud computing

We have also deployed peakScout as a serverless application on Amazon Web Services (AWS), where the analysis environment is containerized and executed via AWS Lambda.Pre-decomposed reference genomes are stored in Amazon S3, retrieved on demand, and decompressed within the execution environment, eliminating the need for persistent infrastructure while supporting multiple genome assemblies. As shown in Figure 2, users interact through a lightweight web client to submit peak files (e.g., BED, narrowPeak) and analysis parameters directly to the AWS Lambda endpoint. Input files are processed against the selected reference genome, and results are returned as compressed artifacts for immediate preview or download. All intermediate files are discarded after execution, ensuring a stateless and reproducible workflow. This cloud-based design enables analyses to complete within seconds of file submission, providing a technically robust yet accessible platform where users simply upload a file, wait several seconds and then retrieve results. The web version of peakScout is available at https://vandydata.github.io/peakScout/.

#### 2.6.3. Use as a command line tool

Several examples for peakScout utilization at the command line are provided in the GitHub README file. In short, one would simply type *peakScout*, followed by the desired function (*peak2gene, gene2peak*, or *decompose_ref*), and finally the required and optional parameters for that analysis. Execution is typically less than 10 seconds when run locally for a peak file containing 50000 peaks. Example output is show in Figure 2C.

## Conclusion

While originally designed for CUT&RUN, CUT&TAG and ChIP-seq, peakScout’s flexible input requirements and support for popular peak caller outputs (MACS2, SEACR) makes it broadly applicable to other genomic methods that generate coordinate-based regions of interest, including ATAC-seq, DNase-seq, and FAIRE-seq experiments. The lack of advanced parameters are intentional to make things easy and accessible for all non-technical users, or those who desire a first pass at results before engaging in deep bioinformatics analysis.

## Acknowledgements

None at this time.

## Data Availability

We have made available prep-processed gene annotation references for common organisms, publicly available and cloud-ready at s3://cds-peakscout-public/. Example peakScout outputs are available as supplementary data 1 (XLSX) and supplementary data 2 (CSV).

